# The reproducibility of research and the misinterpretation of *P* values

**DOI:** 10.1101/144337

**Authors:** David Colquhoun

**Affiliations:** University College London

## Abstract

We wish to answer this question If you observe a “significant” *P* value after doing a single unbiased experiment, what is the probability that your result is a false positive?. The weak evidence provided by *P* values between 0.01 and 0.05 is explored by exact calculations of false positive risks.

When you observe *P* = 0.05, the odds in favour of there being a real effect (given by the likelihood ratio) are about 3:1. This is far weaker evidence than the odds of 19 to 1 that might, wrongly, be inferred from the *P* value. And if you want to limit the false positive risk to 5 %, you would have to assume that you were 87% sure that there was a real effect before the experiment was done.

If you observe *P* =0.001 in a well-powered experiment, it gives a likelihood ratio of almost 100:1 odds on there being a real effect. That would usually be regarded as conclusive, But the false positive risk would still be 8% if the prior probability of a real effect were only 0.1. And, in this case, if you wanted to achieve a false positive risk of 5% you would need to observe *P* = 0.00045.

It is recommended that the terms “significant” and “non-significant” should never be used. Rather, *P* values should be supplemented by specifying the prior probability that would be needed to produce a specified (e.g. 5%) false positive risk. It may also be helpful to specify the minimum false positive risk associated with the observed *P* value.

Despite decades of warnings, many areas of science still insist on labelling a result of *P* < 0.05 as “statistically significant”. This practice must contribute to the lack of reproducibility in some areas of science. This is before you get to the many other well-known problems, like multiple comparisons, lack of randomisation and *P*-hacking. Precise inductive inference is impossible and replication is the only way to be sure,

Science is endangered by statistical misunderstanding, and by senior people who impose perverse incentives on scientists.

## 1. Introduction

> “The major point of this paper is that the test of significance does not provide the information concerning psychological phenomena characteristically attributed to it; and that, furthermore, a great deal of mischief has been associated with its use. What will be said in this paper is hardly original. It is, in a certain sense, what “everybody knows.” To say it “out loud” is, as it were, to assume the role of the child who pointed out that the emperor was really outfitted only in his underwear. Little of that which contained in this paper is not already available in the literature, and the literature will be cited”
>
> Bakan, D. (1966) *Psychological Bulletin*, 66 (6), 423 - 437

When you have done an experiment, you want to know whether you have made a discovery or whether your results could have occurred by chance. More precisely, what you want to know is that when a statistical test of significance comes out positive, what is the probability that you have a false positive i.e. there is no real effect and the results have occurred by chance. This probability is defined here as the *false positive risk* (FPR). In [1] it was called the false discovery rate (FDR), but false positive risk is perhaps a better term because it is almost self-explanatory and because it avoids confusion with the problem of multiple comparisons where the term FDR is commonly used (see also Appendix A1).

The question to be answered is, as before [1], as follows.

If you observe a “significant” *P* value after doing a single unbiased experiment, what is the probability that your result is a false positive?

The experiment is assumed to be randomised and unbiased, with all of the assumptions that were made in calculating the *P* value being exactly true, It is also assumed that we are concerned with a single experiment so there are no problems of multiple comparisons. Real life can only be worse, so in that sense the results given here are the most optimistic possible.

The problem of multiple comparisons is often an important source of false discoveries, but isn’t discussed in this paper. It is worth noting that all the methods for correcting for multiple comparisons aim to correct only the type 1 error. The result is therefore a (corrected) *P* value, so it will still underestimate the false positive risk, for the reasons to be described.

It’s assumed throughout this paper that we wish to test a precise hypothesis, e.g. that the effect size is zero (though it makes little difference if we allow a narrow band around zero [2] [3]). The reasonableness of this approach is justified in Appendix A1.

Most discussions of this topic use the standardised normal distribution (*z* values). But most samples are small, often so small that the experiment is underpowered [4], so here we use the distribution of Student’s *t*.

The discussion will be framed as a comparison between the means of two independent samples, each of *n* normally-distributed observations. The assumptions of Student’s *t* test are therefore fulfilled exactly,

Recently it was asserted that if we observe a *P* value just below 0.05, then there is a chance of at least 26% that your result is a false positive [1]. In that paper attention was concentrated on *P* values that came out close to 0.05, and the results were found by repeated simulations of *t* tests. The aim now is to extend the results to a range of *P* values, and to present programs (in R), and a web calculator, for calculation of false positive risks, rather than finding them by simulation. Better ways of expressing uncertainty are discussed, namely likelihood ratios and reverse Bayesian inference.

It is recommended that the terms “significant” and “non-significant” should never be used. Rather, *P* values and confidence intervals should be supplemented by specifying also the prior probability that would be needed to produce a specified (e.g. 5%) false positive risk.

Before getting to results it will be helpful to clarify the ideas that will be used.

## 2. Definition of terms

A *P* value is defined thus.

> If there were actually no effect (if the true difference between means were zero) then the probability of observing a value for the difference equal to, or greater than, that actually observed is called the *P* value. In other words the *P* value is the chance of seeing a difference at least as big as we have done, if, in fact, there were no real effect.

This definition sounds a bit tortuous, and it’s quite rare for experimenters to be able to define the *P* value accurately. But even when you have the definition right, it’s hard to see exactly what the *P* value tells us. The most common (mis)interpretations are “the *P* value is the probability that your results occurred by chance”. Or “the *P* value is the probability that the null hypothesis is true”. Both of these are disastrously wrong [5]. The latter definition is obviously wrong because the *P* value is calculated on the premise that the null hypothesis is true, so it can’t possibly tell you about the truth of the null hypothesis. The former is wrong because in order to calculate the probability that the result occurred by chance, we need the total number of positive tests, not only those that are found when the null hypothesis is true (see Figure 2 in ref [1]).

The *P* value does exactly what it says. Clearly, the smaller the *P* value, the less likely is the null hypothesis. The problem lies in the fact that *there is no easy way to tell how small P must be* in order to prevent you from making a fool of yourself by claiming that an effect is real when in fact it isn’t. The probability that your results occurred by chance is not the *P* value: it is the false positive risk [5].

The terms used to describe a null hypothesis significance test (NHST), in this case a Student’s *t* test, are defined in Figure 1. The type 1 error rate (in this case 5%) is the probability of finding a “significant” result, *given that the null hypothesis is true*. Because, like the *P* value, it is conditional on the null hypothesis being true, it can’t tell us anything about the probability that the null is true and can’t tell us anything direct about the false positive risk. For that we need to know also what happens when the null hypothesis isn’t true.

In order to calculate the false positive risk the null hypothesis is not enough. We need also an alternative hypothesis. This is needed because, as Berkson said in 1942 [6].

**Figure 1.**
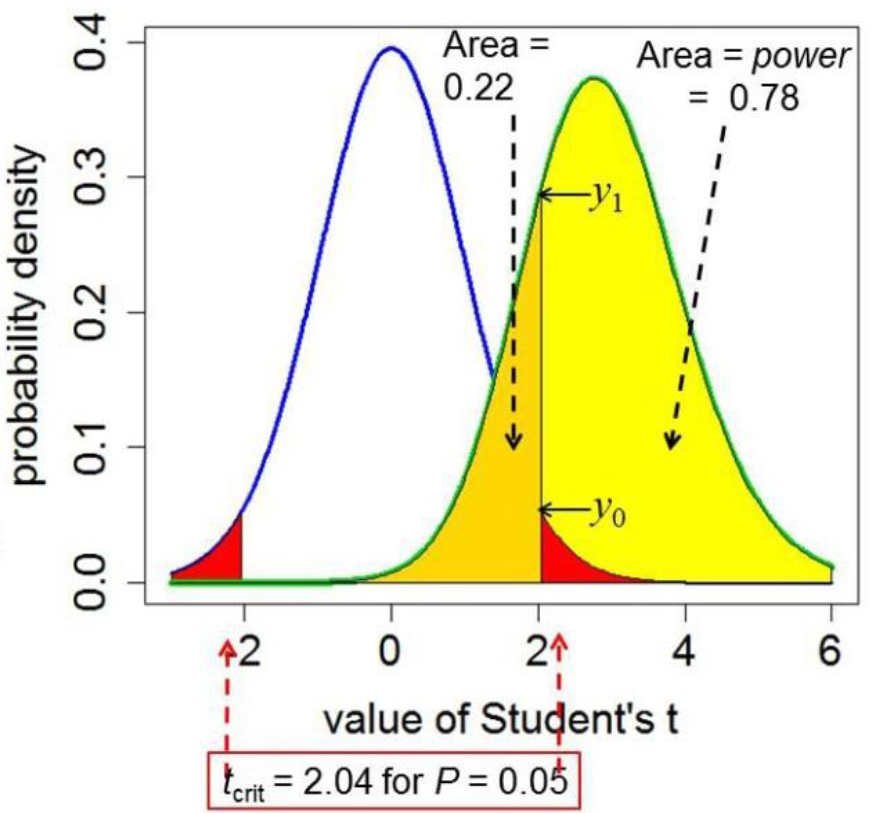
Definitions for a null hypothesis significance test. A Student’s *t* test is used to analyse the difference between the means of two groups of *n* = 16 observations. The *t* value therefore has 2(*n* − 1) = 30 degrees of freedom. The blue line represents the distribution of Student’s *t* under the null hypothesis (H_0_): the true difference between means is zero. The green line shows the noncentral distribution of Student’s *t* under the alternative hypothesis (H_1_): the true difference between means is 1 (one standard deviation). The critical value of *t* for 30 df and *P* = 0.05 is 2.04, so, for a two-sided test, any value of *t* above 2.04, or below–2.04, would be deemed “significant”. These values are represented by the red areas. When the alternative hypothesis is true (green line), the probability that the value of *t* is below the critical level (2.04) is 22% (gold shaded are): these represent false negative results. Consequently, the area under the green curve above *t* = 2.04 (shaded yellow) is the probability that a “significant” result will be found when there is in fact a real effect (H_1_ is true): this is the *power* of the test, in this case 78%. The ordinates marked *y*_0_ (= 0.526) and. *y*_1_ (= 0.290) are used to calculate likelihood ratios, as in section 5.

> “If an event has occurred, the definitive question is not, “Is this an event which would be rare if null hypothesis is true?” but “Is there an alternative hypothesis under which the event would be relatively frequent?”

Or, paraphrasing Sellke *et al.* 2001 [7]]

> “knowing that the data are ‘rare’ when there is no true difference is of little use unless one determines whether or not they are also ‘rare’ when there is a true difference”.

The quantities defined in Figure 1 are not sufficient to define the false positive risk. To get what we want we need Bayes’ theorem. It can be written thus.

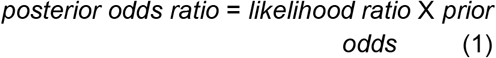

(see eq, A8 for a more precise definition, and appendix in ref [1]), The word ‘prior’ signifies ‘before the experiment;, and ‘posterior’ signifies after the experiment. So the likelihood ratio measures the evidence provided by the experiment. Often we shall prefer to speak of probabilities rather than odds. The probability that a hypothesis is true is related to the odds ratio in favour of the hypothesis being true, thus

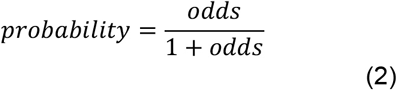

or, conversely,

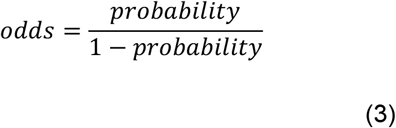

For example, if the odds on the null hypothesis being true were 9 times greater than the odds of the alternative hypothesis being true (odds ratio of 9 in favour of the null hypothesis) then the probability of the null hypothesis is 9/((9 +1)=0.9, and the probability of the null being false is 1 – 0.9 = 0.1.

The prior probability of an event means the probability before the experiment was done. In the context of diagnostic screening tests, it’s well defined, as the prevalence of the condition in the whole population being tested (see Figure 1 in ref [1]). The perpetual warfare over the use of Bayes’ theorem in the context of tests of significance stems from the fact that we hardly ever know a value for this prior probability, and that problem will be discussed later.

Before getting to that, we need to clarify an important distinction,

## 3. Which interpretation is better: ‘*p-less-than*’, or ‘*p-equals*’?

This is an old, but often neglected question. . Although this discussion has gone on for decades in the statistical literature (eg ref [22]), it is unknown to most users. It was discussed in section 10 of ref [1].

The question arises when we try to answer the question: what is the probability that our results occurred by chance if, in a single unbiased test, we find *P* = 0.047 (or whatever value is observed); Is it appropriate to consider all tests that produce *P* ≤ 0.047 or should we consider only tests that give *P* = 0.047? Let’s call these, respectively, the “*p-less-than*” interpretation and the “*p-equals*” interpretations.

The distinction sounds subtle, but simulations make its meaning obvious. Suppose sets of 100,000 *t* tests are simulated, as In ref. [1]. The simulations are intended to mimic what’s done in real life, so each set of simulated data is analysed with a two-independent-sample *t* test (the only difference from real life is that the simulated data are known to fulfil exactly the assumptions made by the test). Each simulated test generates a *P* value. Simulated data are generated for the case when the null hypothesis is true, and separately for the case when the null hypothesis is not true. Thus, unlike in real life, one knows, for each test, whether or not the null hypothesis was true: this makes it possible to count how often the null hypothesis is rejected wrongly and hence the false positive risk can be estimated (see Figure 2 in ref [1]). The calculation of each *P* value involves tail areas in the standard way, i.e. it takes into account all results that depart from the null hypothesis by as much as, *or more than*, the observed amount. But, having generated 100,000 *P* values, there are two possible ways to analyse them. We can look at all of the tests that give *P* values that are equal to *or less than* the observed value (0.047, for example). Or one can look at only the tests that result in *P* values that come out close to 0.047, as observed.

- The *p-equals* interpretation counts the fraction of false positives among all tests that come out with *P* values equal to the observed one, e.g. 0.047.
- The *p-less-than* interpretation counts the fraction of false positives among all tests that come with *P* equal to *or less than* the observed value.

In order to answer our question, we have to regard the outcome of our actual experiment as being a random instance from the 100,000 possible outcomes that were simulated. Since our actual experiment came out with *P* = 0.047, we are interested in simulated experiments that produce *P* values close to 0.047. In other words, the *p-equals* case is what we need to answer our question,

In our one actual experiment there is a fixed true effect size and the prior probability that there’s a real effect is also fixed, though its value is unknown. It makes no sense to select at random, from a prior distribution, a different value of the true effect size for each simulated *t* test (see Appendix A1). The idea of an experiment being a random instance from a large number of imaginary repetitions of the same experiment is a standard way of looking at inference,

Although the distinction between the *p-less-than* case and the *p-equals* case is most easily understood by simulations, one aim of this paper is to supply code that calculates the *p-equals* case exactly (rather than the *p-close-to* case that is all that can be done by simulation), as explained in Appendix A2.-

Since the outcome of the experiment, in our example, was *P* = 0.047 (or whatever value is observed), it seems clear that the ‘*p-equals*’ case is appropriate for interpretation of our particular experiment. Recall that we are not trying to calculate our lifetime false positive risk, but just trying to interpret our single result. Simulations that came out with a *P* value of less than 0.047 were not observed in the real experiment, so they are irrelevant. Most papers (e.g. Wacholder, 2004 [8] and Ioannidis, 2005 [9]) consider only the ‘*p-less-than*’ case, which is easy to calculate, but which, in my opinion, answers the wrong question.

## 4. Simulation *versus* exact calculation

In ref [1], the problem was solved by simulation. For the example used there, we calculated the difference between the means of two groups, each with *n* observations. The difference between the two means can be called the effect size. If the two groups were given treatments that were equally effective, the effect size would be zero *on average*. To simulate experiments in which the null hypothesis is true we generate random samples of *n* “observations” from the same normal distribution e.g. both samples come from a normal distribution with a mean of zero.

In order to calculate the FPR we need to postulate an alternative to the null hypothesis. Let’s say that the true effect size is equal to one, the same as the standard deviation of the individual responses. This is not as arbitrary as it seems at first sight, because identical results are obtained with any other true effect size, as long as the sample size is adjusted to keep the power of the test unchanged [10].

In order to simulate an experiment in which the null hypothesis is not true we generate random samples of *n* ‘observations’ from a normal distribution with a mean of zero for one sample, and for the other sample we take *n* observations from a normal distribution with a mean of one. Both distributions have the same true standard deviation, equal to one.

For example, with a true effect size of one standard deviation, the power of the test is 0.78, for *P* = 0.05, when the sample sizes are *n* = 16.

For each pair of samples, a standard Student’s *t* test is done. Notice that this is an ideal case, because it’s known that the assumptions of the *t* test are obeyed exactly. Real life can only be worse.

It seems beyond doubt that the ‘*p* equals’ case is what we need. Our real experiment came out with *P* = 0.047 (or whatever), so what we need to do is to look at the false positive risk for experiments that produce *P* values of 0.047. If, as in [1], this is done by simulation, one has to look at a narrow band of *P* values around the observed value, say *P* values that lie between 0.045 and 0.05, in order to get adequate numbers of simulated *P* values that are close to the observed value, 0.047 (or whatever). In this paper we calculate exactly the false positive risk that corresponds to any specified *P* value that we have found in a real experiment. An R script is provided to do this calculation (*calc-FPR+LR.R*), and a web calculator [44]. The calculation is outlined in Appendix A2.

The script and web calculator both give also the false positive risk for the ‘*p-less-than*’ case, though this can be found from the tree diagram approach, or calculated simply from equation A4 in Colquhoun (2014) [1].

The difference between the two approaches is illustrated in Figure 2. This shows the false positive risk plotted against the *P* value. The plots are for a well-powered experiment. The curves are calculated with *n* = 16 observations in each sample, because this gives a power of 0.78 for *P* = 0.05 and the specified effect size and standard deviation. The sample size is fixed because it is good practice to estimate sample size in advance to give adequate power at a specified *P* value, usually 0.05.

The top row in Figure 2 is calculated on the basis that the probability that our experiment would have a real effect was 0.1 before the experiment was done: this prior probability shows some scepticism about whether a real effect exists. It might, for example, be appropriate when testing a putative drug, because most drugs candidates fail.

The bottom row in Figure 2 was calculated on the premise that there is is a prior probability 0.5 that our experiment truly had a real effect (a 50:50 chance). This is the largest prior that can legitimately be assumed, in the absence of good empirical data to the contrary (see Figure 5).

**Figure 2.**
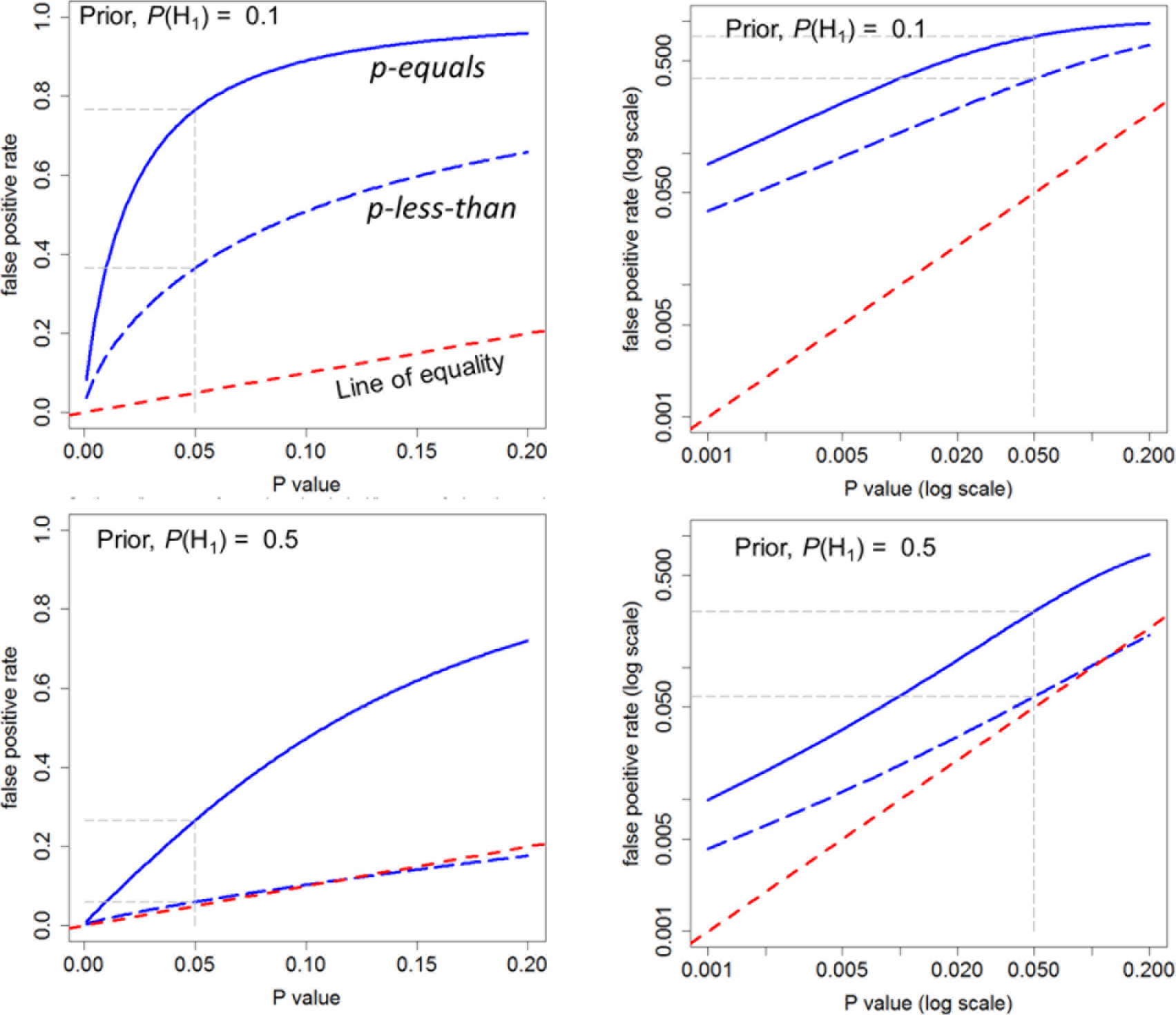
Plots of false positive rate (FPR) against *P* value, for two different ways of calculating FPR. The continuous blue line shows the *p-equals* interpretation and the dashed blue line shows the *p-less-than* interpretation. These curves are calculated for a well-powered experiment with a sample size of *n* = 16. This gives power = 0.78, for *P* = 0.05 in our example (true effect = 1 SD). Top row. Prior probability of a real effect = 0.1 Bottom row. Prior probability of a real effect = 0.5 The dashed red line shows a unit slope: this shows the relationship that would hold if the FPR were the same as the *P* value. The graphs In the right hand column are the same as those in the left hand column, but in the form of a log-log plot. Graphs produced by *Plot-FPR-vs-Pval.R* [44]

The dashed red line in each graph shows where the points would lie if the FPR were the same as the *P* value (as is commonly, but mistakenly, supposed). It’s clear that the FPR is always bigger, often much bigger, than the *P* value over the whole range.

Not surprisingly, the FPR is always bigger when calculated by the *p-equals* method than it is when calculated with the *p-less-than* method. For a *P* value close to 0.05, and prior probability of a real effect = 0.5, the FPR is 26% according to the *p-equals* interpretation, in agreement with the simulations in [1], but the FPR is only 6% according to the *p-less-than* interpretation. When the prior probability of a real effect is only 0.1, the FPR for a *P* value of 0.05 is 76% for the *p-equals* interpretation (again agreeing with the value found by simulation in [1]). But according to the *p-less-than* interpretation the FPR is 36% (in agreement with the tree diagram approach and the calculation in appendix A4 of.ref. [1]).

It’s clear from Figure 2 that the *only* case in which the FPR is similar to the *P* value is when the prior probability of a real effect is 0.5 and we use the inappropriate *p-less-than* interpretation. In this case, the bottom row in Figure 2 shows that the FPR (dashed blue line) is only just above the *P* value for *P* values close to 0.05, though for *P* = 0.001 the FPR is 5-fold greater than the *P* value, even in this case‥ But, as discussed above, the appropriate answer to the question is given by the *p-equals* interpretation, and the fact that this suggests a false positive risk of 26% for an observed *P* value close to 0.05 was what led to the conclusion in [1] that the false positive risk is *at least* 26% and for an implausible hypothesis (with a low prior probability) it will be much higher.

## 5. Likelihood ratios

It has often been suggested that it would be better to cite likelihood ratios rather than *P* values, e.g. by Goodman [11],[12] [39].

The word likelihood is being used here in a particular statistical sense. The likelihood of a hypothesis is defined as a number that is directly proportional to the probability of observing the data, given a hypothesis. Notice that this is *not* the same thing as the somewhat elusive probability of the hypothesis given the data: that is harder to find. The calculation of the likelihood is entirely deductive (under our assumptions-see Appendix A1), so it doesn’t involve induction: see ref [5]). When we speak of a maximum likelihood estimate of a parameter it means that we choose the value that makes our observations more probable than any other.

The likelihood of a hypothesis is not interpretable on its own: we can interpret only the relative likelihood of two hypotheses. This is called the likelihood ratio. In the example used here (and in ref [1]), the two hypotheses are the null hypothesis (the true difference between means is zero) and the alternative hypothesis (the true difference between means is one).

The likelihood ratio is the part of Bayes’ theorem (eq. 1) that describes the evidence provided by the experiment itself. And it is the part of Bayes’ theorem that can be calculated by two methods, the *p-equals* method and the *p-less-than* method (see section 3).

Notice that use of the likelihood ratio avoids the problem of deciding on the prior probabilities. That, at least, is true when we are testing precise hypotheses (see Appendix A1).

Suppose, for example that the data are such that a *t* test gives *P* = 0.05. The probability of observing *P* = 0.05 exactly under the null hypothesis is proportional to the ordinate labelled *y*_0_ in Figure 1, and the probability of observing *P* = 0.05 exactly under the alternative hypothesis is proportional to the ordinate labelled *y*_1_ in Figure 1.

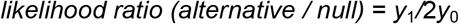

The factor of 2 arises because we are talking about a 2-sided test. This is discussed in more detail in Appendix A2.

Values for likelihood ratios are printed out by the R script, *calc-FPR+LR.R*, and are given by the web calculator [44]. Part of the output file from the script is shown in Table 1 and Table 2.

If we observe *P* = 0.05, for which *power* = 0.78, as in Figure 1, the likelihood ratio for the alternative versus the null is 2.76 (see Table 1 and Appendix A2 for details). So the alternative hypothesis is only 2.76 times as likely as the null hypothesis (not 20 times as likely as might, wrongly, be guessed from the observed *P* value, 0.05). This is one reason for thinking that the *P* value (as usually misinterpreted) exaggerates the strength of the evidence against the null hypothesis [11],[12].

There are two ways to calculate the likelihood ratio. The method just described is the *p-equals* interpretation (see section 3 and Fig 2). This is the appropriate way to answer our question. We can also calculate the likelihood ratio in a way that’s appropriate for the *p-less-than* interpretation. In this case the likelihood ratio is simply equal to the relative areas under the curves in Figure 1, i.e. *power/P value.* In the example in Fig 1, this is 0.78/0.05 = 15.6 e., the alternative hypothesis is 15.6 times as likely as the null hypothesis. This calculation was done in the appendix in ref [1], but it is not appropriate for answering our question.

**Table 1.**
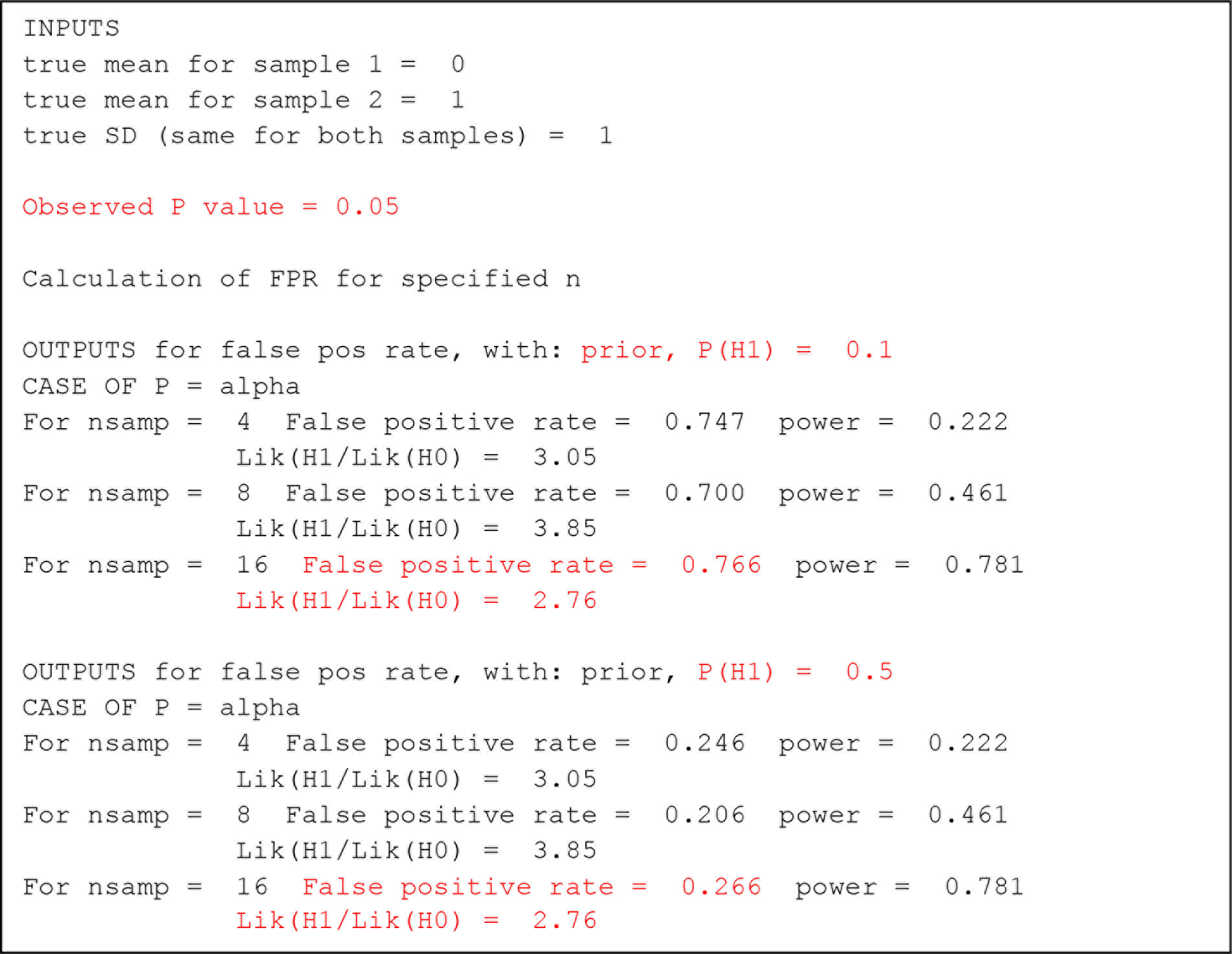
The case when we observe *P* = 0.05. Sample output from the R script, *calc-FPR+LR.R*. Values mentioned in the text in red.

The fact that we hardly ever have a valid value for the prior probability means that it’s impossible to calculate the false positive risk. Therefore rigorous induction is impossible [5]. But we can give a minimum value for the FPR.

*Observed likelihood ratios*. The likelihood ratios just discussed were calculated for the true effect size (of 1 SD). This is not known in real life. So we may ask what happens if we calculate the likelihood ratio from our experimental data? This is easily answered by simulation. Rather than calculating the likelihood ratio for a specified constant effect size (1 SD) and a specified standard deviation, we calculate separately for each simulated experiment the likelihood ratio for the ‘observed’ effect size, sample standard deviation and the *P* value. This is done using the R script *two_sample-simulation-+LR+prior.R* (see ref [44]).

The likelihood ratios, of course, vary from one simulated experiment to the next, but if we look only at experiments that come out with *P* values close to 0.05, say 0.0475 < *P* < 0.0525, the likelihood ratios (in favour of there being a real effect) for these are all close to 3.64. This is a bit bigger than the theoretical value of 2.76, and that is to be expected because it’s calculated for each simulated experiment using the observed effect size and the observed effect size is, in this case, the maximum likelihood estimate of the true effect size. But the odds of there being a real effect are still much smaller than the 19:1 odds that might, wrongly, be inferred from the *P* value of 0.05.

If these simulations are repeated for *P* values that are close to 0.01 (looking only at simulated experiments that come out with 0.0095 < *P* < 0.0105) we find that the likelihood ratio in favour of there being a real effect is 15.4 (and in this case the theoretical value is much the same). So observation of *P* = 0.01 makes the alternative hypothesis (a real effect) 15.4 times more likely than the null hypothesis (no real effect). This makes the existence of a real effect much less likely that the 99 to 1 odds that might, wrongly, be inferred from observing a *P* value of 0.01. In fact it doesn’t even reach the common standard of 19 to 1.

The likelihood ratio is the bit of Bayes’ theorem that tells us about the evidence from the experiment. The fact that observing *P* = 0.05 corresponds with a likelihood ratio of only about 3 in favour of the alternative hypothesis is a good reason to be sceptical about claiming that there’s a real effect when you observe a *P* value close to 0.05. It also shows that the *P* value is a very imperfect measure of the strength of the evidence provided by the experiment.

However calculating the likelihood ratio still doesn’t tell us what we really want to know, the false positive risk. Just as there is no easy way to tell how small a *P* value must be to provide reasonable protection against false positives, so there is also no easy way to know how big the likelihood ratio (in favour of there being a real effect) must be to provide reasonable protection against false positives. What we really want to know is the false positive risk, and for that we need a Bayesian approach.

Notice that Bayes’ theorem (eq. 1) states that when the prior odds are 1 (*i.e.* the prior probability of there being a real effect is 0.5) the posterior odds are equal to the likelihood ratio. So the likelihood ratio does give us a direct measure of the *minimum* false positive risk (given that it’s usually not acceptable to assume any higher prior than 0.5). In this case, the theoretical likelihood ratio, when we observe *P* = 0.05 in our experiment, is 2.76. Thus the posterior odds in favour of there being a real effect is, in this case, also 2.76. The posterior probability of there being a real effect is 2.76/(2.76 + 1) = 0.734, from eq. 2. And therefore the posterior probability that the null hypothesis is true, the false positive risk, is 1-0.734 = 0.266. The minimum false positive risk is thus 26%, as found by calculation above and by simulation in [1]. The likelihood ratio found from the experimental results (which is what you can calculate in practice), was slightly bigger, 3.64, so this implies a minimum false positive risk of 1– (3.64/(1 + 3.64)) = 21.6 %, slightly better, but not by much.

If we observe in our experiment that *P* = 0.01, the false positive risk will be lower. In this case the likelihood ratio in favour of there being a real effect is 15.4. In the most optimistic case (prior probability for a real effect of 0.5) this will be the posterior odds of there being a real effect. Therefore, the posterior probability of the null hypothesis, as above, 1– (15.4/(15.4 + 1)) = 0.061, far bigger than the observed *P* value of 0.01. It doesn’t even reach the usual 5% value.

These values are all minimum false positive risks. If the prior probability of a real effect is smaller than 0.5, the false positive risk will be bigger than these values, as shown in Figure 5.

## 6. False positive risk as function of sample size

The R programs, or the web calculator [44], make it easy to calculate the FPR for any given *P* value, with different sample sizes. The calculation is outlined in Appendix A2. Figure 3 shows such graphs for sample sizes of *n* = 4, 8 and 16, as used in [1]. These sample sizes give the power of the *t* tests, at the *P* = 0.05 point, as 0.78 (*n* = 16), 0.46 (*n* = 8) and 0.22 (*n* = 4). These values cover the range of powers that are common in published work [4].

The FPR is calculated by the *p-equals* method (see section 3 and Figure 2), using the R script *Plot-FPR-vs-Pval.R,* see ref [44]). The program produces also graphs calculated by the *p-less-than* interpretation, but this isn’t what we need to answer our question.

As in Figure 2, the dashed red line shows where the points would lie if the FPR were equal to the *P* value. The right hand column shows a log-log plot of the graph in the left hand column. It’s obvious that in all cases, the false positive risk is a great deal bigger than the *P* value.

The top row of graphs in Figure 3 is calculated with a prior probability that there is a real effect of 0.1, i.e. the existence of a real effect is somewhat implausible. For a *P* value close to 0.05, the FPR is 76% for the well powered sample size (*n* = 16, power = 0.78), as found by simulation in [1], and by calculation: see Table 1.

The lower row of graphs in Figure 3 is calculated assuming a prior probability of a real effect of 0.5. In other words, before the experiment is done there is assumed to be a 50:50 chance that there is a real effect so the prior odds are 1. This is usually the largest prior probability that can reasonably be assumed (see discussion and Figure 5). For the well-powered experiment (*n* = 16, power = 0.78) the FPR is 26% when a *P* value of 0.05 is observed (see Table 1), Again this agrees with the value found by simulation in [1].

The graphs in Figure 3 show also that the curves for different sample sizes are quite close to each other near *P* = 0.05. This explains why it was found in ref [1] that the FPR for *P*=0.05 was insensitive to the power of the experiment. The fact that the FPR can actually be slightly lower with a small sample than with a big one is a well-understood phenomenon: see, for example, Senn (Chapter 13 in ref. [13]), and Holder (2009) [40].

**Figure 3.**
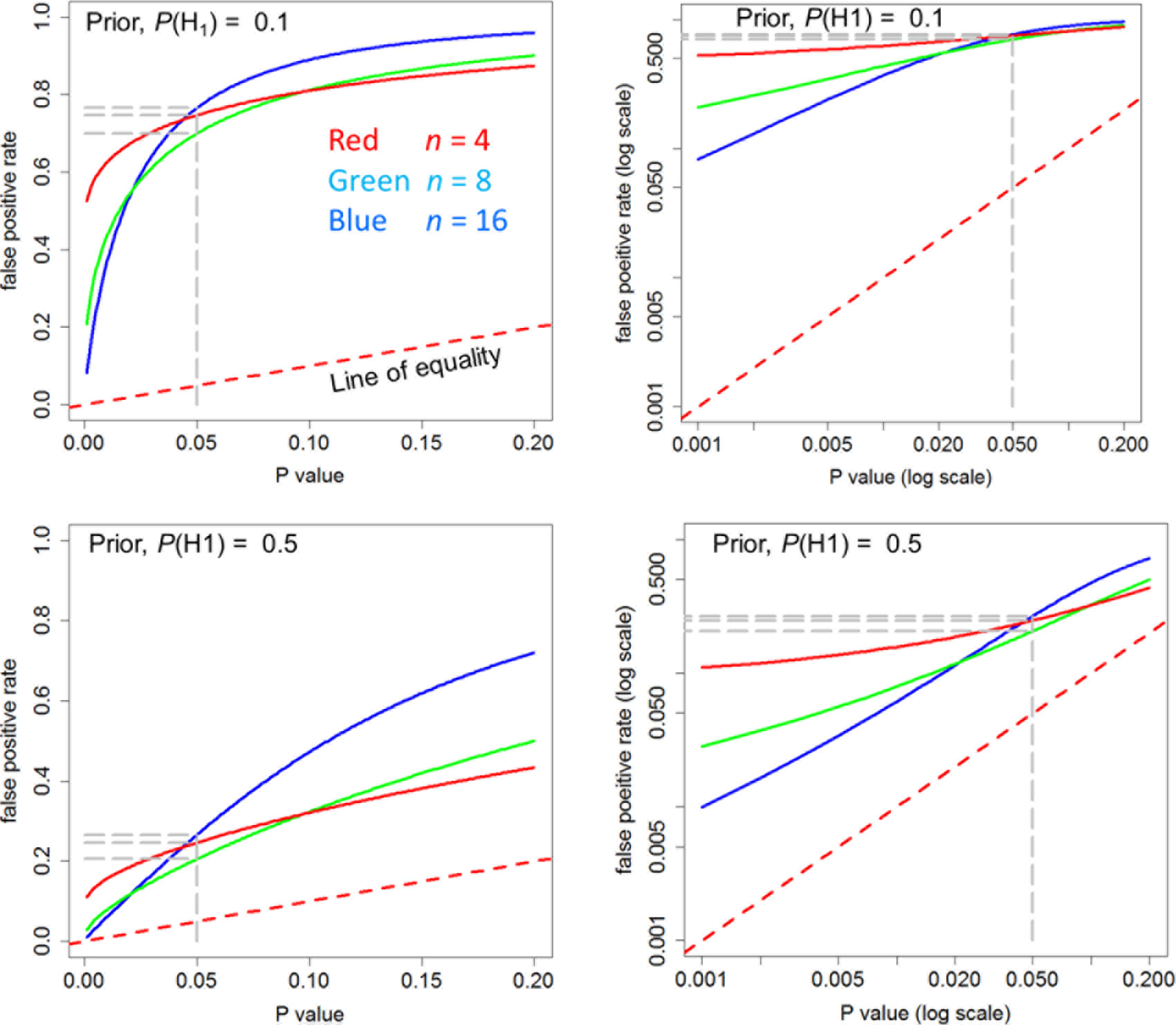
The calculated false positive rate plotted against the observed *P* value. The plots are for three different sample sizes: *n* = 4 (red), *n* = 8 (green) and *n* = 16 (blue), Top row. Prior probability of a real effect = 0.1 Bottom row. Prior probability of a real effect = 0.5 The dashed red line shows a unit slope: this shows the relationship that would hold if the FPR were the same as the P value. The graphs In the right hand column are the same as those in the left hand column, but in the form of a log-log plot. Graphs produced by *Plot-FPR-vs-Pval.R* [44]

For smaller observed *P* values, Figure 3 shows that in all cases the false positive risk is much greater than the observed *P* value.

For example, if we observe a *P* value of 0.001, we can see what to expect by running the R script *calc-FPR+LR.R,* with the observed *P* value set to 0.001 (see Table 2). It can also be calculated with the web calculator [44], as shown in Figure 4. These values give a likelihood ratio of 100 to 1 in favour of there being a real effect. If we assume that the prior probability of a real effect is 0.5 then this corresponds to a minimum false positive risk of 1.0%. That is 10 times the *P* value but stillprovides good evidence against the null hypothesis.

However with a prior probability of 0.1 for a real effect (an implausible hypothesis), as in Figure 4, the false positive risk is still 8%, despite having observed *P*=0.001. It would not be safe to reject the null hypothesis in this case, despite the very low *P* value and the large likelihood ratio in favour of there being a real effect.

**Figure 4.**
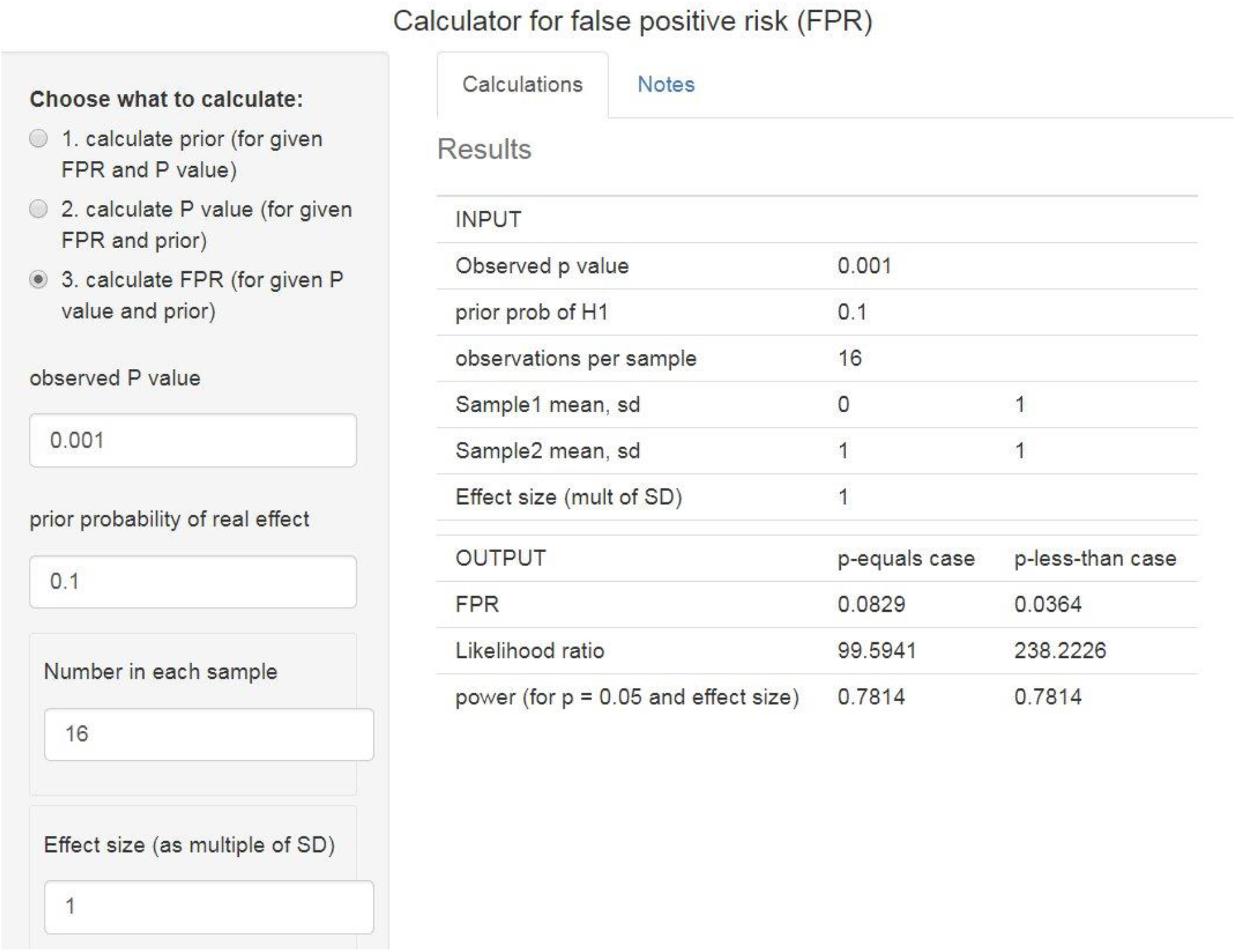
Web calculator [44] for the case where we observe a *P* value of 0.001 and the prior probability of a real effect is 0.1. http://fpr-calc.ucl.ac.uk/

An alternative way to look at the problem is to specify a false positive risk that you find acceptable, and to calculate the *P* value that would be needed to achieve it. This can be done with the R script *calc_p-val.R,* or with the web calculator [44]. If we are willing to make fools of ourselves 1 in 20 times, we would specify a false positive risk of 5%. With a well-powered experiment (*n* = 16), to achieve a false positive risk of 0.05 we would need a *P* value of *P* = 0.0079 if the prior probability of a real effect were 0.5 (the most optimistic case). But if the prior probability of a real effect were only 0.1, we would need to observe *P* = 0.00045.

These examples serve to show that it would be foolish to ignore the prior probability, even though we don’t know its value.

**Table 2.**
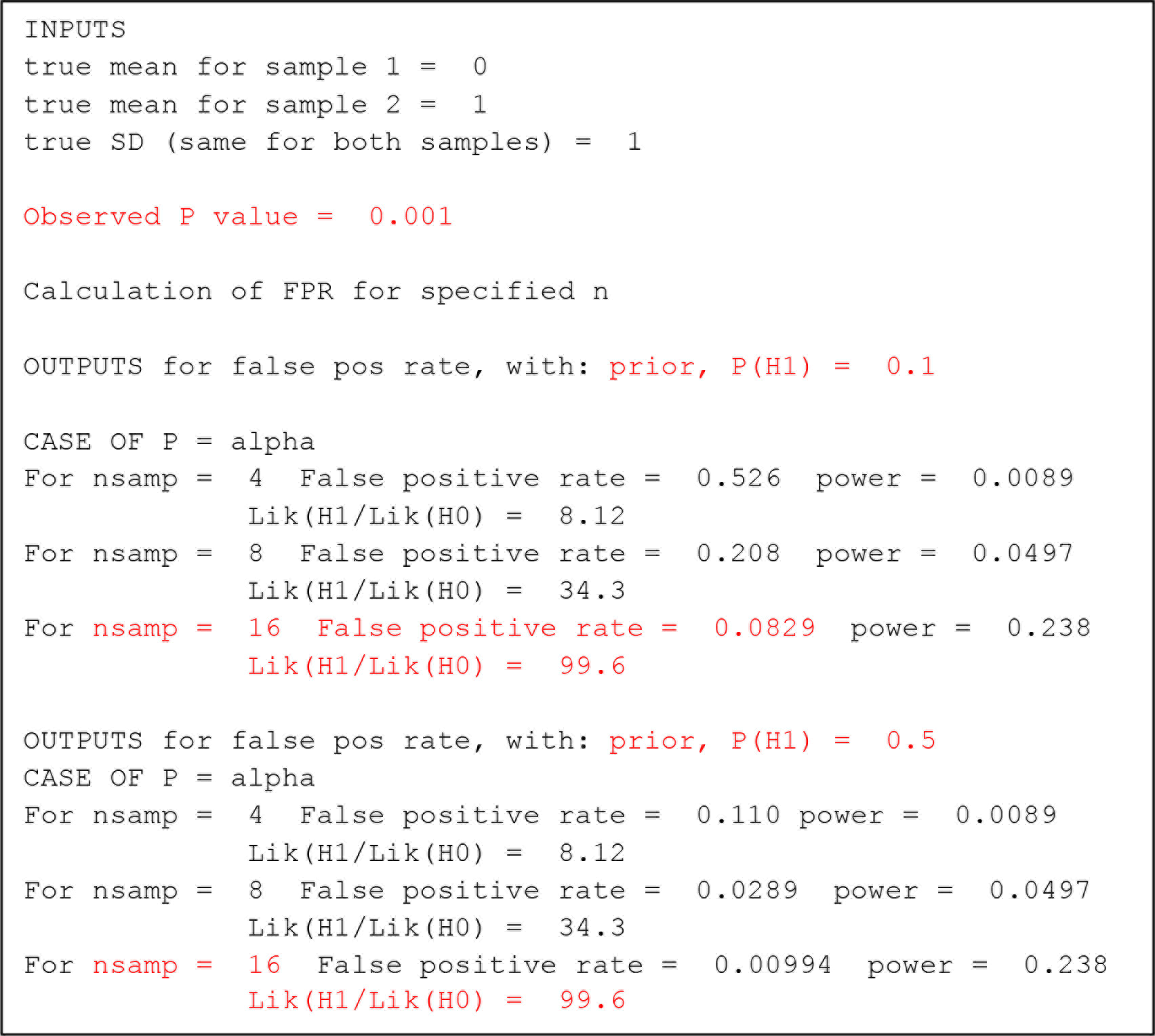
The case when we observe *P* = 0.001. Sample output from the R script, *calc-FPR+LR.R* [44]. Values mentioned in the text in red.

Figure 5 shows how the false positive risk varies with the prior probability of there being a real effect. It’s calculated for a well-powered experiment (0.78) that gives a *P* value just below 0.05 (see legend for details).

**Figure 5.**
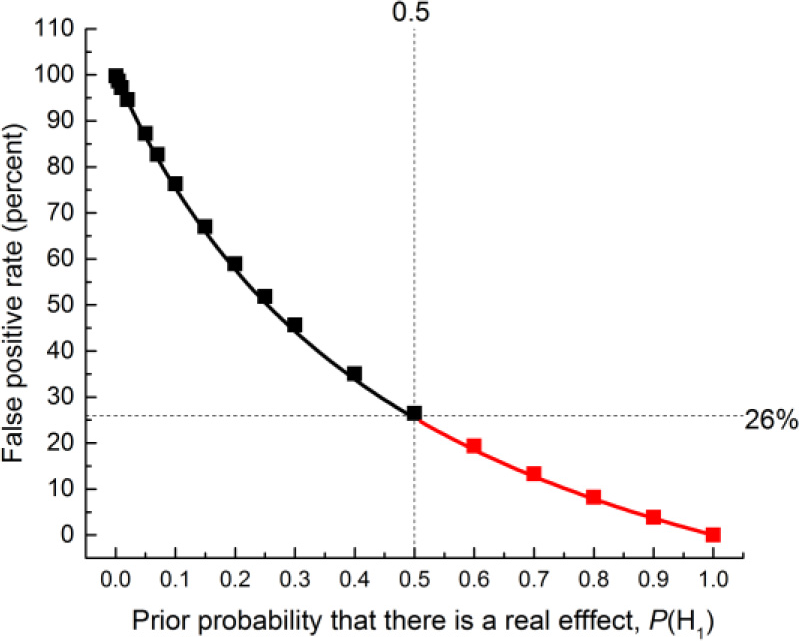
The false positive rate plotted against the prior probability for a test that comes out with a *P* value just below 0.05. The points for prior probabilities greater than 0.5 are red because it is essentially never legitimate to assume a prior bigger than 0.5. The calculations are done with a sample size of 16, giving power=0.78 for *P*=0.0475. The square symbols were found by simulation of 100,000 tests and looking only at tests that give *P* values between 0.045 and 0.05. The fraction of these tests for which the null hypothesis is true is the false positive rate. The continuous line is the theoretical calculation of the same thing: the numbers were calculated with *origin-graph.R* and transferred to Origin to make the plot.

As stated before, the false positive risk is 26% for a prior 0.5, but for a less plausible hypothesis, with a prior probability of 0.1, the false positive risk is 76%. If the same treatment were given to both groups (or, equivalently, a dummy pill was given to one group, and a homeopathic pill was given to the other) then the prior probability is zero: in this case the null hypothesis is true, so 5% of tests come out positive but what matters is that the false positive risk is 100%. At the other extreme, if we were totally sure that there was a real effect before doing the experiment (prior probability = 1) then all positive tests would be true positives and the false positive risk would be zero.

The folly of ignoring the prior probability can also be illustrated starkly by an example based on decision-making in a court of law [14]. In the “Island Problem” discussed there, the probability of observing the evidence, given that the suspect was guilty was 0.996 (see ref [14] for details). But that alone tells us nothing about what we need to know, namely the probability that the suspect is guilty, given the evidence. To mistake the former for the latter is the error of the transposed conditional [5], or, in this context, the prosecutor’s fallacy. So would it be more helpful to calculate the likelihood ratio as an indication of the strength of the evidence? This is Prob(*evidence* | *guilty*) / Prob(*evidence* | *not guilty*) which evaluates to odds of 250:1 that a suspect was guilty rather than not guilty, in the light of the evidence. That sounds beyond reasonable doubt. But in that (somewhat artificial) example, the prior odds of guilt were known to be 1 in 1000, so, from Bayes’ theorem (equation 1), the posterior odds of guilt are not 250:1, but rather 0.25:1. In other words there are odds of 4 to 1 *against* guilt. Use of the likelihood ratio alone would probably have led to a wrongful conviction (and, in some countries, execution) of an innocent person. See ref [14] for details.

The prior probability of there being a real effect (or, in this example, the prior probability of guilt) may not be known, but it certainly can’t be ignored. Luckily there is a solution to this dilemma. It will be discussed next,

## 7. The reverse Bayesian argument

In all the examples given so far, it has been necessary to specify a prior probability in order to calculate the FPR. But we hardly ever have a valid value for this prior. Matthews [15] has proposed an ingenious way round the problem posed by the inconvenient fact that we essentially never have a valid value for the prior probability that there is a non-zero effect. He suggests that we reverse the argument. We specify a false positive risk that’s acceptable to us, and calculate the prior probability that would be needed to achieve that rate. We can then judge whether or not that prior probability is, or is not, plausible. The calculation is outlined in Appendix A3. The calculations are done by the R script, *calc-prior.R,* or with the web calculator [44].

Similar proposals have been made by others, especially Held (2013) [41].

For example, if we observe a *P* value close to 0.05, and we want a false positive risk of 5% (which is what many people mistakenly think the *P* value gives you), that implies that you must assume that the prior probability of a non-zero effect is 87% (for sample size *n*= 16). In other words, to obtain a false positive risk of 5% you have to be almost sure (prior=0.87) that there is a non-zero effect *before* doing the experiment. The web calculator for this case is shown in Figure 6.

And In order to get a false positive risk of 1% we would have to assume a prior probability of 0.98. These priors are obviously preposterously high. It is yet another way of looking at the weakness of the evidence provided by a *P* value close to 0.05.

**Figure 6.**
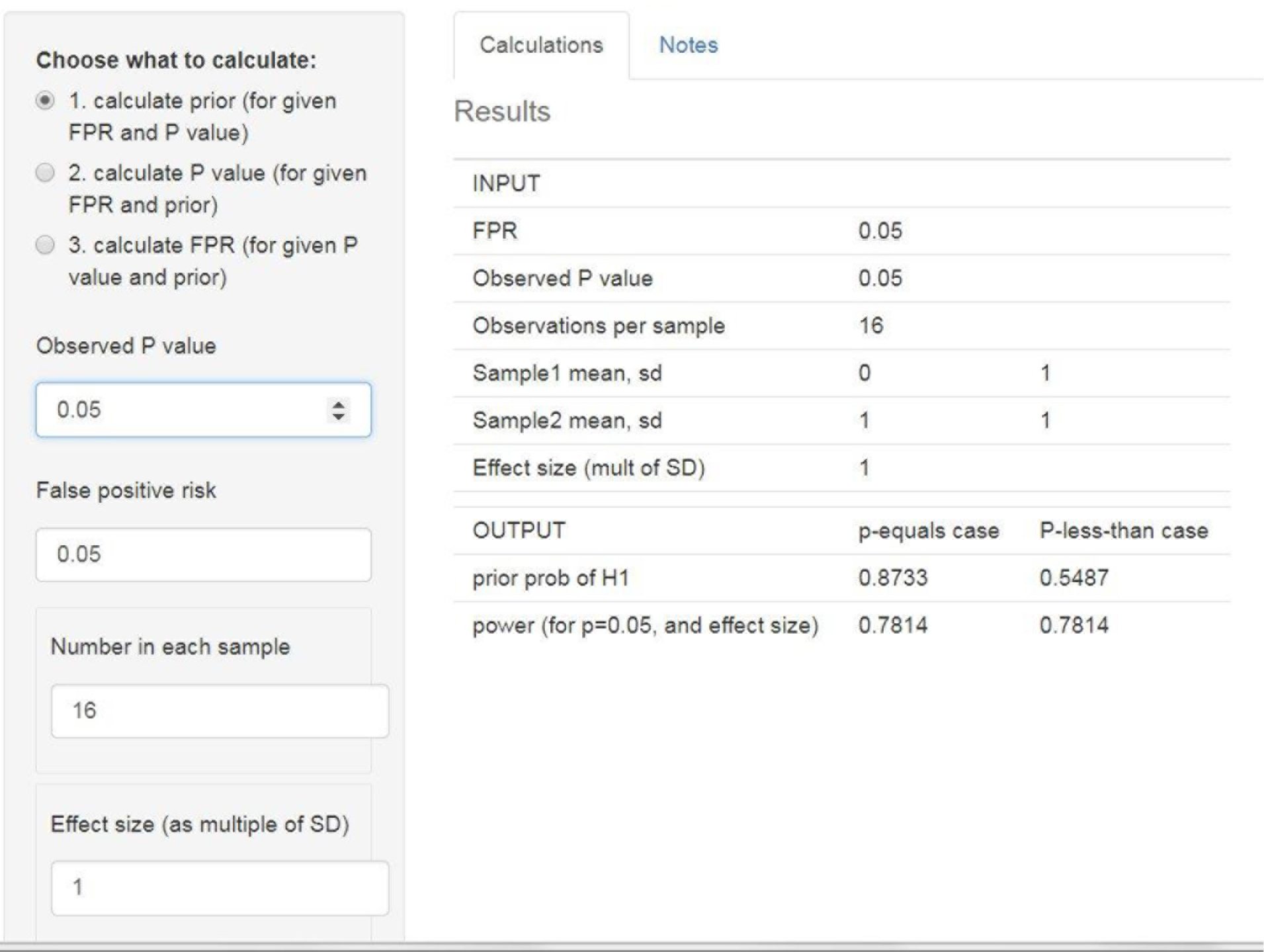
Web calculator[44]: calculation of the prior probability than would be needed to achieve a false positive risk of 5% when we observe *P* = 0.05. http://fpr-calc.ucl.ac.uk/

If we observe a *P* value close to 0.01, then to achieve a false positive risk of 5% we would have to assume a prior probability that there is a real effect of 0.55, i.e. that before the experiment was done, it was (slightly) more probable than not that there was a real effect. And to achieve a false positive risk of 1%, the prior would have to be 0.87, unacceptably high.

If we observed a *P* value of 0.001, then to achieve a false positive risk of 5%, we’d have to assume the prior was 0.16. That’s not impossible insofar as it’s below 0.5, but if the hypothesis were implausible (e.g. we were testing homeopathic pills) it might still be thought implausibly high. A false positive risk of 0.01 (ten times larger than the *P* value) would need a prior of 0.50: you then have to decide whether or not it’s reasonable to assume that, before you have the data, there’s a 50:50 chance that there’s a real effect.

These priors are calculated using the true mean difference and true standard deviation, and, in real life, these are not known. As in the case of the likelihood ratio, we may ask what happens if we calculate the prior probability from our experimental data? Again this is easily answered by simulation. Rather than calculating the prior probability for a specified constant effect size (1 SD) and a specified standard deviation, we calculate separately for each simulated experiment the prior probability using the ‘observed’ effect size, sample standard deviation and *P* value. This is done using the R script *two_sample-simulation-+LR+prior.R* (see ref [44]). This gives very similar results to the exact calculation. For example, the prior probability that we need to postulate in order to achieve a false positive risk of 5% is close to 0.84 for ‘experiments’ that come out with *P* values close to 0.05 (between *P=*0.0475 and 0.0525), for sample size *n*=16. This is close to the prior of 0.087 found with the true effect size and SD. For smaller *P* values the difference is even smaller. It will therefore be quite good enough to calculate the prior probability from the observed effect size and standard deviation (e.g using ***calc-prior.R*, or the web calculator [44]**).

Of course the judgement of whether or not the calculated prior probability is acceptable or not is subjective. Since rigorous inductive inference is impossible [5], some subjective element is inevitable. Calculation of the prior probability that is needed to achieve a specified false positive risk is a lot more informative than the equally subjective judgement that *P* < 0.05 is adequate grounds for claiming a discovery.

## 8. Discussion

The fact that we hardly ever have a valid value for the prior probability means that it’s impossible to calculate the false positive risk. Therefore rigorous induction is impossible [5].

Although it is often said that *P* values exaggerate the evidence against the null hypothesis, this is not strictly true. What *is* true is that *P* values are often misinterpreted as providing more evidence against the null hypothesis than is the case. Despite the fact that the smaller the *P* value, the less plausible is the null hypothesis, there is no simple way to know how small the *P* value must be in order to protect you from the risk of making a fool of yourself by claiming that an effect is real when in fact the null hypothesis is true, so all you are seeing is random sampling error.

The misconception that a *P* value is the probability that your results occurred by chance, i.e. that it’s the probability that the null hypothesis is true, is firmly lodged in the minds of many experimenters. But it is wrong, and it’s seriously misleading.

Table 3 shows a summary of some of the results. Notice particularly the strong effect of the prior probability. If it were possible to assume that a real effect were as likely as not (prior probability 0.5) then an observation of *P* = 0.005 would imply a reasonable false positive risk of 3.4%. But if the prior probability were only 0.1, then *P* = 0.005 would give a disastrous FPR of 24%. Even *P*=0.001 would give FPR of 8% in this case. To reduce the FPR to 5% you would need *P* = 0.00045

**Table 3.**
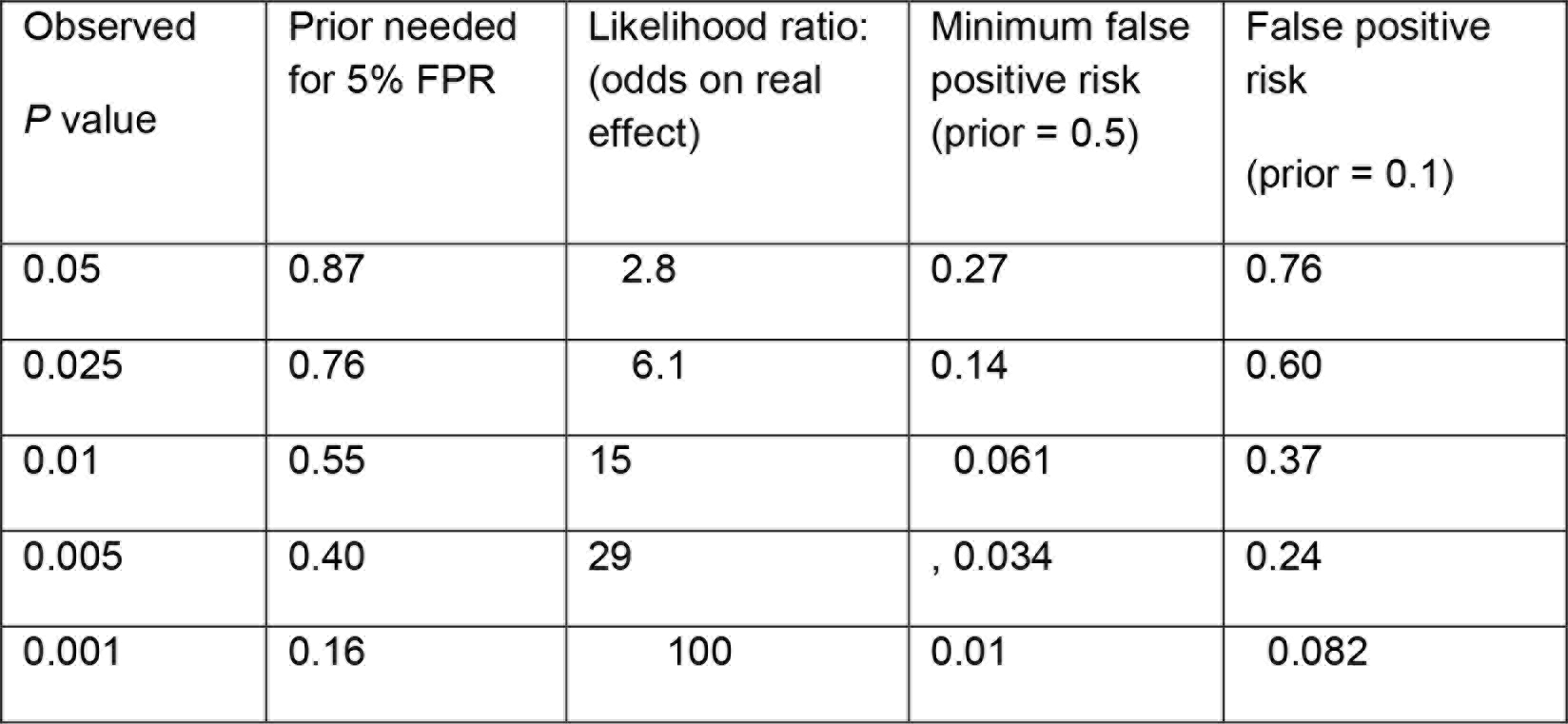
Summary of results (calculated by the *p-equals* method, for a well-powered experiment, *n* = 16. Calculations done with *calc-prior.R* and ***calc-FPR+LR.R*** [44]). Column 1: the observed *P* value Column 2: the prior probability of there being a real effect that it would be necessary to postulate in order to achieve a false positive rate (FPR) of 5% (see section 7) Column 3: likelihood ratios (likelihood for there being a real effect divided by likelihood of null hypothesis, See section 5 and appendix A2) Column 4: the minimum false positive rate, i.e. the FPR that corresponds to prior odds of 1 (see Figure 5). Column 5. the false positive rate that would be expected if the prior probability of there being a real effect were only 0.1

| Observed<br>$P$ value | Prior needed<br>for 5% FPR | Likelihood ratio:<br>(odds on real<br>effect) | Minimum false<br>positive risk<br>(prior = 0.5) | False positive<br>risk<br>(prior = 0.1) |
| --- | --- | --- | --- | --- |
| 0.05 | 0.87 | 2.8 | 0.27 | 0.76 |
| 0.025 | 0.76 | 6.1 | 0.14 | 0.60 |
| 0.01 | 0.55 | 15 | 0.061 | 0.37 |
| 0.005 | 0.40 | 29 | 0.034 | 0.24 |
| 0.001 | 0.16 | 100 | 0.01 | 0.082 |

The problem of reproducibility has not been helped by the inability of statisticians to agree among each other about the principles of inference. This is shown very clearly by the rather pallid nature of the statement about *P* values that was made by the American Statistical Association [16]. It said what you shouldn’t do, but failed to say what you should do, Reading the 20 accompanying statements shows little sign of convergence among the views of the warring camps. Stephen Senn put it thus in a tweet.

@stephensenn Replying to @david_colquhoun @david_colquhoun You know us statisticians at the moment we are struggling to converge on whether we are converging or not.

The inability to agree is also made clear by the Royal Statistical Society discussion about the ASA statement [17], and by Matthews’ assessment of it, one year later [18].

Even such gurus of evidence-based medicine as Heneghan and Goldacre don’t mention the contribution made by the myth of *P* values to the unreliability of clinical trials [19]

More surprisingly, even some accounts of significance testing by professional statisticians don’t always point out the weakness of *P* values as evidence. Their teaching to biologists must bear some of the blame for widespread misunderstanding.

Despite the controversy that still surrounds the Bayesian approach, it’s clear that we are all Bayesians at heart. This is illustrated by the aphorisms “extraordinary claims require extraordinary evidence”, and “if sounds too good to be true, it’s probably untrue”. The problems arise when we want to put numbers on the uncertainty. And the main problem is the impossibility of putting numbers on the prior probability that the null hypothesis wrong (see Appendix A1).

A real Bayesian would specify a prior distribution, which, they would claim, represents the state of knowledge before the experiment was done, based on current expert opinion. This appears to be nothing more than an appeal to authority (see Edwards (1992) [20]. There is essentially never enough expert opinion to specify a prior distribution, and to try to do so carries the risk of reinforcing current prejudices. The result will depend on which expert you ask. There will be as many different answers as there are experts. That is not helpful: in fact it’s fantasy science. So what can we do?

The way to get round the problem proposed by Colquhoun (2014) [1] was to say that any prior probability greater than 0.5 is unacceptable because it would amount to saying that you know the answer before the experiment was done. So you can calculate a false positive risk for a prior probability of 0.5 and describe it as a *minimum false positive risk*. If the hypothesis were implausible, the prior probability might be much lower than 0.5, and the false positive risk accordingly much higher than the minimum. But setting a lower bound on the false positive risk is a lot better than ignoring the problem.

Using likelihood ratios in place of *P* values has been advocated, e.g. [11],[12]. They have the advantages that (under our assumptions-see Appendix A1) they can be calculated without specifying a prior probability, and that they are the part of Bayes’ theorem (eq. 1) that quantifies the evidence provided by the experiment (something that *P* values don’t do [20]).

Calculation of likelihood ratios certainly serves to emphasise the weakness of the evidence provided by *P* values (section 5, and [21]): if you observe *P* = 0.05, the likelihood ratio in favour of there being a real effect is around 3 (section 5), and this is pretty weak evidence, Even if we observe *P* = 0.01, the likelihood of there being a real effect is only about 15 times greater than the likelihood of the null hypothesis. So the existence of a real effect is much less likely that the 99 to 1 odds that might be, wrongly, inferred from the observed *P* value of 0.01. In fact it doesn’t even reach the common standard of 19 to 1.

Useful though likelihood ratios can be, they aren’t a solution to the problem of false positives, for two reasons (see section 5). Firstly, there is no simple way to tell how big the likelihood ratio must be to prevent you from making a fool of yourself. And secondly, likelihood ratios can overestimate seriously the strength of the evidence for there being a real effect when the prior probability is small. Their use could result in conviction of an innocent person (section 5). Even if we observe *P*=0.001, which gives a likelihood ratio of 100 in favour of there being a real effect, the false positive risk would still be 8% if the prior probability of a real effect were only 0.1 (see Table 2).

I suggest that the best way of avoiding the dilemma posed by the unknown prior is to reverse the argument and to calculate, using the observed *P* value, what the prior probability would need to be to achieve a specified false positive risk (section 7, and ref [15]). This can be calculated easily with the web calculator [44], for which it is the default option (see Figure 6). This procedure leaves one with the subjective judgement of whether or not the calculated prior is reasonable or not (though if the prior comes out bigger than 0.5, it is never reasonable, in the absence of hard evidence about the prior distribution).

If we observe a *P* value close to 0.05, then, in order to achieve a false positive risk of 5% it would be necessary to assume that the prior probability that there was a real effect would be as high as 0.87. That would be highly unreasonable.

Other ways to get round the problem of the unknown prior have been proposed. A full Bayesian analysis involves choosing a prior distribution for the effect size. It has been shown that it’s possible to specify a prior distribution that gives the biggest probability of rejecting the null hypothesis [7] [22] [23]. Even these priors, when we observe a *P* value close to 0.05, give a posterior probability of the null hypothesis being true of over 20% (i.e. the odds on the null being false are less than 4 to 1). That is far weaker evidence against the null hypothesis than the (wrongly-interpreted) *P*=0.05 might suggest. These mathematically-sophisticated Bayesian arguments lead to very similar conclusions to those given here. For example, when *P*=0.05 and prior probability is 0.5, Berger *et al.* [7] [22] find a false positive risk of at least 22% and Johnson [23] finds 17-25%. Both are close to the value of 26% found here by simpler arguments.

## 9. Conclusions: and what should be done?

One thing that you should *not* do is to follow the advice given by many journals: Some statement like the following is only too common [24].

> “a level of probability (*P*) deemed to constitute the threshold for statistical significance should be defined in Methods, and not varied later in Results (by presentation of multiple levels of significance). Thus, ordinarily *P* < 0.05 should be used throughout a paper to denote statistically significant differences between groups.”

As Goodman [11] said

> “The obligatory statement in most research articles, ‘p values below 0.05 were considered statistically significant’ is an empty exercise in semantics”

Not only is *P* < 0.05 very weak evidence for rejecting the null hypothesis, but statements like this perpetuate the division of results into “significant” and “non-significant”.

At the other extreme, neither should you use a fantasy prior distribution to do a full Bayesian analysis. Valen Johnson has said, rightly, that

> “subjective Bayesian testing procedures have not been—and will likely never be— generally accepted by the scientific community” [25].

So here is what I think should be done.

1. Continue to give *P* values and confidence intervals. These numbers should be given because they are familiar and easy to calculate, not because they are very helpful in preventing you from making a fool of yourself. They do not provide good evidence for or against the null hypothesis. Giving confidence intervals has the benefit of focussing attention on the effect size. But it must be made clear that there is *not* a 95% chance that the true value lies within the confidence limits you find. Confidence limits give the same sort of evidence against the null hypothesis as *P* values, i.e, not much.
2. I propose that the best way of indicating the strength of evidence provided by a single *P* value is to use the reverse Bayesian method (section 7). That is, calculate what prior probability would be needed to achieve a specified false positive risk (eg. use *calc-prior.R,* or the web calculator [44]). If this prior is bigger than 0.5, then you aren’t on safe ground if you claim to have discovered a real effect. If the calculated prior is less than 0.5 then It is then up to you to argue that the calculated prior is plausible, and up to the reader to judge your whether or not they are convinced by your argument. For example, *P* = 0.005 would require a prior of 0.4 in order to achieve a 5% false positive risk (see Table 3). So, if you observe *P* = 0.005. and you are happy with a 5% false positive risk, it’s up to you, and the reader, to judge whether or not the prior of 0.4 is reasonable or not. This judgement is largely subjective, and people will disagree about it. But inference has to involve subjectivity somewhere. Calculation of the prior seems to me to be a better way than specifying an arbitrary prior in order to calculate an FPR. In the end, only replication will resolve arguments.
3. Perhaps most important of all, never, ever, use the words “significant” and “non-significant” to describe the results. This wholly arbitrary dichotomy has done untold mischief to the integrity of science.
4. A compromise idea is to change the words used to describe observed *P* values. It has been suggested that the criterion for statistical significance be changed from 0.05 to 0.005 [23] [39]. Certainly, one should never use the present nonsensical descriptions: *P* > 0.05 not significant; *P* < 0.05 significant; *P* < 0.01 very significant. And one should never use asterisks do denote them. Reduction of the threshold for “statistical significance” to *P* =0.005 would certainly reduce the number of false positives, but there is no such thing as a threshold. And of course a threshold of *P* = 0.005 would result in missing many real effects. In practice, decisions must depend on the relative costs (in money and in reputation) that are incurred by wrongly claiming a real effect when there is none, and by failing to detect a real effect when there is one. Marginal *P* values are fine as a signal to investigate further. In the end, the only solution is replication.
5. Another way of looking at the strength of the evidence provided by a single *P* value is to state, as well as the *P* value, the likelihood ratio or, better, the corresponding minimum false positive risk (e.g. use *calc-FPR+LR.R* or the web calculator [44]). These are much better ways of assessing the evidence provided by the experiment than simply stating a *P* value‥
6. Always be aware that no method exists for rigorous inductive argument [5]. In practice, judgement, and especially replication, is always needed. There is no computer program that can automatically make a judgement for you. Here’s a real life example. A study of transcranial electromagnetic stimulation, published In *Science*, concluded that it “improved associative memory performance”, *P* = 0.043 [26]. If we assume that the experiment had adequate power (the sample size of 8 suggests that might be optimistic) then, in order to achieve a false positive risk of 5% when we observe *P* = 0.043, we would have to assume a prior probability of 0.85 that the effect on memory was genuine (found from *calc-prior.R*, or the web calculator [44]). Most people would think it was less than convincing to present an analysis based on the assumption that you were almost certain (probability 0.85) to be right before you did the experiment. Another way to express the strength of the evidence provided by *P*=0.043 is to note that it makes the existence of a real effect only 3.3 times as likely as the existence of no effect (likelihood ratio found by *calc-FPR+LR.R* or the web calculator [44]). This would correspond to a minimum false positive risk of 23% if we were willing to assume that nonspecific electrical zapping of the brain was as likely as not to improve memory (prior odds of a real effect was 1). Recently a paper (with 72 authors) has appeared [39] which proposes to change the norm for “statistical significance” from *P* = 0.05 to *P* = 0.005. Benjamin *et al.* [39] makes many of the same points that are made here, and in [1]. But there are a few points of disagreement.

1. Benjamin *et al.* propose changing the threshold for “statistical significance”, whereas I propose dropping the term “statistically significant” altogether: just give the *P* value and the prior needed to give a specified false positive risk of 5% (or whatever). Or, alternatively, give the *P* value and the minimum false positive risk (assuming prior odds of 1). Use of fixed thresholds has done much mischief.
2. The definition of false positive risk in equation 2 of Benjamin *et al.* [39] is based on the *p-less-than* interpretation. In [1], and in this paper, I argue that the *p-equals* interpretation is more appropriate for interpretation of single tests. If this is accepted, the problem with *P* values is even greater than stated by Benjamin *et al.* (e.g see Figure 2).
3. The value of *P* =0.005 proposed by Benjamin *et al.* [39] would, in order to achieve a false positive risk of 5%, require a prior probability of real effect of about 0.4 (from *calc-prior.R*, or the web calculator, with power = 0.78, i.e. *n* =16). It is, therefore, safe only for plausible hypotheses. If the prior probability were only 0.1, the false positive risk with *P* = 0.005 would be 24% (from *calc-FPR+LR.R,* or the web calculator, with *n* = 16). It would still be unacceptably high even with *P* =0.005. Notice that this conclusion differs from that of Benjamin et al [39] who state that the *P* = 0.005 threshold, with prior = 0.1, would reduce the false positive risk to 5% (rather than 24%). This is because they use the *p-less-than* interpretation which, in my opinion, is not the correct way to look at the problem.

Many reported *P* values fall in the marginal range, between 0.01 and 0.05 [27,28]. They provide only weak evidence against the null hypothesis. This suggests that the problem of false positives is likely to be responsible for a substantial part of the reported lack of reproducibility. Although the problems outlined here have been known to statisticians for at least 70 years, they are still largely unknown to experimenters.

It is hard to avoid the conclusion that experimenters don’t want to know about the myth of *P* < 0.05. Despite the decades for which statisticians have been pointing out the inadequacies of this approach, practice has hardly changed. Indeed it is still widely used in papers that have professional statisticians as co-authors. Experimenters perceive that to abandon the myth of *P* < 0.05 might harm their place in the academic rat race.

Journals must bear some of the blame too. Their statistical advice is mostly more-or-less inaccurate. But when I pointed out the harm that would be done by such bad advice [24], the response of journal editors was to say that if they were to adopt recommendations of the sort given above it would “damage their journals’ impact factors”. The effect of competition between journals is as corrupting as the effect of competition between individuals.

This paper addresses a very limited question: how do you interpret the result of a single unbiased test of significance. It makes no attempt to estimate the science-wide false positive rate. The increased awareness if the problem of reproducibility has led to many attempts to assess the scale of the problem. Since most of these papers use the *p-less-than* approach to calculate false positive risks, the problem may be even worse than they suggest. See, for example [9] [29] [30] [33] [42] [43].

The people who must bear the ultimate responsibility for this sad state of affairs are university presidents and heads of research funders. While they continue to assess “productivity” by counting publications, counting citations and judging papers by the journal in which they are published, the corruption will continue. Despite abundant evidence that metrics such as these don’t measure quality, and do encourage bad practice, they continue to be widespread [29] [30] [32] [42] [43]. .The efforts of university administrators to edge up a place or two in university rankings can be very cruel to individuals, even leading to death [30], and the people who do this seem to have no appreciation of the fact that the rankings with which they are so obsessed are statistically illiterate [31–33].

One really disastrous aspect of the rash of false positives is that it gives ammunition to people who distrust science. Until recently, these were largely homeopaths and other such advocates of evidence-free medicine. Now, with a president of the USA who denies the effectiveness of vaccination, and doubts the reality of climate change, the need for reliable science is greater than it ever has been.

It has become a matter of urgency that universities, politicians and journals should stop subverting efforts to improve reproducibility by imposing perverse incentives on the people who do the work. These pressures have sometimes led to young scientists being pushed by their seniors into behaving unethically. They are fighting back [38] and that’s a good augury for the future

## Appendix

## A1. The point null hypothesis

I’m aware that ‘I’m stepping into a minefield and that not everyone will agree with the details of my conclusions. However I think that most statisticians will agree with the broad conclusion that *P* values close to 0.05 provide weak evidence for a real effect and that the way in which *P* values continue to be misused has made a substantial contribution to Irreproducibility.

It should be pointed out that the conclusions here are dependent on the assumption that we are testing a point null hypothesis. The null hypothesis is that the true effect size is zero and the alternative hypothesis is (for a twosided test) that the true effect size is not zero. In other words the prior distribution for the null hypothesis is a spike located at zero effect size [11] [3]. This is, of course, what statisticians have been teaching for almost acentury. This seems to me to be an entirely reasonable approach. We want to know whether our experiment is consistent with a true effect size of zero. We are not asserting that the effect is exactly zero, but calculating what would happen if it were zero. If we decide that our results are not consistent with a true effect size of zero then we can go ahead and calculate our estimate of the effect size. At that point, we then have to decide whether the effect size is big enough to matter. It’s not uncommon in clinical trials to find small effect sizes which, even if they are real, aren’t big enough for a patient to get a noticeable benefit. In basic science, focussing research efforts on small effects with moderate *P* values is undoubtedly driving extensive investigations of epiphenomena that cost a great deal of money and ultimately serve no-one.

It is not uncommon to hear the view that the point null hypothesis is not worth testing because it is never exactly true. Apart from the fact that it can be exactly true (just give the same pill to both groups), this seems irrelevant because it makes little difference if we allow a narrow band of values close to zero [3]. What we are asking is whether our observations are consistent with an effect size of zero (or close to zero). This is precisely the question which experimenters want to answer, in order to avoid making fools of themselves by claiming an effect when none exists.

Bayesians may object to this approach and say instead that a prior distribution should be postulated. Senn [34] has pointed out that, if we assume a smeared prior distribution for the null hypothesis, it’s possible to find such a distribution that makes the FPR the same as the *P* value, at least for one-sided tests. In terms of simulations, this means that in each simulated *t* test, we choose at random from this prior distribution, a different value for the null hypothesis. This procedure makes some sense if you are considering a lifetime of experiments. But even in this case, the prior distribution will be different for each sort of experiment, and it will never be known anyway.

In this paper we are considering the interpretation of a single experiment. It has a single true effect size, which we don’t know but wish to estimate. The simulations, here and in ref. [1], assume the same true effect size in every simulation. The only variability is in the “observations” that are used to estimate the effect size‥ We simply count the number of “significant” results that are false positives. It is alarmingly high.

In earlier drafts of this paper, the probability of getting a “significant” *P* value when the null hypothesis is true was dubbed the false positive rate. In ref [1] the same quantity was called the false discovery rate. But since we are talking about the result of a single experiment, the term “false positive risk” is a better description. Others call the same thing the False Positive Report Probability (FPRP), e.g. ref [41]. Unfortunately the nomenclature in this field is a mess, so it’s important, when reading any paper to check the definitions

A noted in the introduction, the problem of multiple comparisons isn’t discussed here. But notice that the methods for correcting for multiple comparisons all aim to correct only the type 1 error: the result is a (corrected) *P* value, As such it is still subject to all the reservations about interpretation of *P* values that are discussed in this paper. In particular, it will still underestimate the false positive risk

## A2 Calculation of likelihood ratios and false positive risks

In ref [1], likelihood ratios and false positive risks were calculated in appendix A2-A4, according to the *p-less-than* interpretation. Here we do analogous calculations for the *p-equals* case. That is what is needed to answer our question (see section 3).

The critical value of the *t* statistic is calculated as

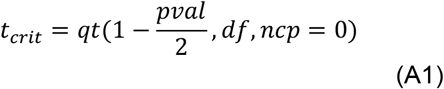

where *qt*() is the inverse cumulative distribution of Student’s *t* statistic. The arguments are the observed *P* value, *pval,* the number of degrees of freedom, *df*, and the non-centrality parameter, *ncp* (zero under the null hypothesis) For the example in Figure 1, *pval* = 0.05, with 30 degrees of freedom so this gives *t*_crit_ = 2.04. Under the null hypothesis (blue line in Figure 1) the probability density that corresponds to the observed *P* value is

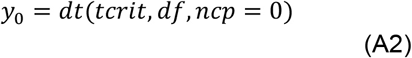

where *dt*() is the probability density function of Student’s *t* distribution with degrees of freedom *df* and non-centrality parameter = zero under the null hypothesis. This is the value marked *y*_0_ in Figure 1 and in that example its value is 0.0526.

Under the alternative hypothesis, we need to use the non-central *t* distribution (green line in Figure 1). The non-centrality parameter is

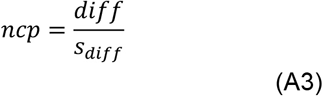

where *diff* is the difference between means (1 in this case) and *s*_diff_ is the standard deviation of the difference between means, 0.354 in Figure 1, so *ncp* = 2.828. The probability density that corresponds with *t*_crit_ = 2.04 is

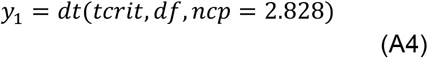

In the example in Figure 1, this is 0.290, labelled *y*_1_ in Figure 1.

The likelihood of a hypothesis is defined as a number that is directly proportional to the probability of making the observations when the hypothesis is true. For the *p-equals* case we want the probability that the value of *t* is equal to that observed. This is proportional to the probability density that corresponds to the observed value (see Figure 1). (More formally, the probability is the area of a narrow band centred on the observed value, but we can let the width of this band go to zero.) For a twosided test, under the null hypothesis the probability occurs twice, once at *t* = –2.04 and once at + 2.04 (see Figure 1). For brevity, let’s define

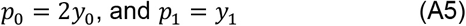

Thus, under the *p-equals* interpretation, the likelihood ratio in favour of H_1_, is

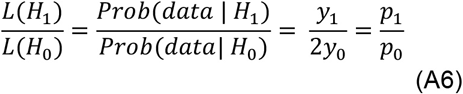

In forming the likelihood ratio, the arbitrary constant in the likelihoods cancels out.

In the example in Figure 1, the likelihood ratio is 0.290/(2 × 0.0526) = 2.76. The alternative hypothesis is 2.76 times more likely than the null hypothesis. This is weak evidence for the existence of a real effect.

In order to calculate the false positive risk, we need Bayes’ theorem. This can be written in the form

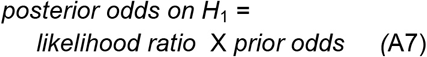

Or, in symbols,

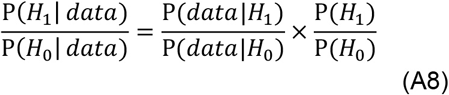

The likelihood ratio represents the evidence provided by the experiment. It is what converts the prior odds to the posterior odds. If the prior odds are 1, so the prior probability of a real effect is 0.5, then the posterior odds are equal to the likelihood ratio. In the example in Figure 1, this is 2.76, so the posterior probability of there being a real effect, from eq. 2, is 2.76/(2.76 +1) = 0.734. And the posterior probability of the null hypothesis is 1 – 0.734 = 0.266. In the case of prior odds = 1 this can be written as

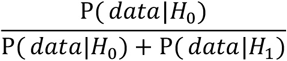

So when we observe *P* = 0.05 there is a 26.6% chance that the null hypothesis is true. And this is the minimum false positive risk of 26.6% that was found above, and by simulations in section 10 of ref [1].

Likelihood ratios tell us about the minimum false positive risk that is found when the prior odds are 1 *i.e*. prior probability = 0.5 (see Figure 5). But to assume this is dangerous (see section 6). In general, we can’t assume that the prior odds are 1. If the prior probability of a real effect is *P*(H_1_), and the prior probability of the null hypothesis is therefore *P*(H_0_) = 1 - *P*(H_1_), then the false positive risk is

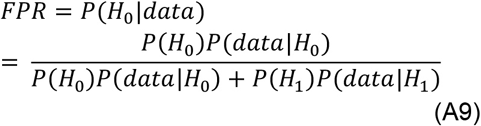

where *data* are represented by the observed *P* value.

In the example given in Figure 1, *P*(*data* | H_0_) = *p*_0_ =2 × 0.0526 = 0.105, and *P*(*data* | H1) = *p*_1_ = 0.290.

If the prior probability of a real effect is *P*(H_1_) = 0.1, so *P*(H_0_) = 0.9, then, for the example in Figure 1, the observed *P* value of 0.05 implies a false positive risk of

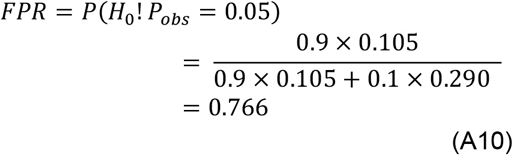

This agrees with the 76% false positive risk found by calculation above, and by simulations in [1].

## A3 The calculation of the prior probability: the reverse Bayesian approach

In order to do this, all we have to do is to rearrange eq A8 to give the prior needed for a specified FPR.

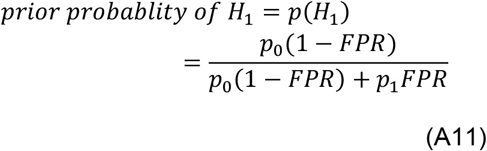

For the example in Figure 1, *p*_0_ = 0.105 and *p*_1_ = 0.290, so if we observe *P* = 0.05, and want a false positive risk of 5%, it would be necessary to assume a prior probability of there being a real effect of

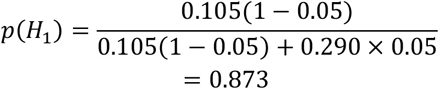

To attain a false positive risk of 5% after observing *P* = 0.05, we would have to be almost (87%) certain that there was a real effect before the experiment was done. Values of the prior needed to give any specified FPR, given an observed *P* value can be calculated using the web calculator (see Figure 6) or by the R script ***calc-prior.R* [44]**.

## A4 How to do the calculations

In practice the calculations will be done with the R scripts or the web calculator [44]. The inputs needed for these calculations will be specified next.

There are three variables, the observed *P* value, the false positive risk (FPR) and the prior probability that the null hypothesis is false. The R scripts or the web calculator will work out any one of these, given numbers for the other two.

All three calculations require also the number of observations in each sample, and the effect size, expressed as a multiple of the standard deviation of the observations (default value 1.0) The default number per sample is 16 which gives a power of 0.78 for P = 0.05 and effect size = 1-see ref [1] and [3] for more details.

In order to test the null hypothesis, *H*_0_, that the true effect size is zero we have to specify an alternative hypothesis, *H*_1_, which is that the true effect size is not zero. But note that all that matters is the effect size expressed as a multiple of the standard deviation of the original observations (sometimes known as *Cohen’s d*). The true mean of sample 1 is always 0 (null hypothesis), The true mean of sample 2 is set to the normalised effect size so the true standard deviation can always be set to 1, with no loss of generality.

Exactly the same results will be found for any other effect size as long as the sample size is adjusted to keep the power constant. For example effect size = 1 and *n* = 16 gives power = 0.78. For an effect size of 0.5 SD, *n* =61 gives similar power and similar FPR etc. And for an effect size of 0.2 SD, a power of 0.78 requires *n* = 375, and again this gives similar FPR etc‥ So choose *n* so that the calculated power matches that of your experiment [10].

## A5. Bayesian estimation in single molecule kinetics

It has become common to see Bayesian arguments in many fields, including my erstwhile job, interpretation of single molecule kinetics. Experimental data in that field come as distributions (of sojourn times in the open and shut states of a single ion channel), so maximum likelihood estimation was the obvious thing to do, once the mathematical problems had been solved [35,36]. Recently the same problem has been approached by Bayesian methods [37]. The parameters to be estimated are the rate constants for transitions between discrete states of the receptor protein. The priors were taken to be uniform between lower and upper limits. The limits are dictated by physical considerations, and the same limits were applied in maximum likelihood as constraints on the parameter values. Under these circumstances, the point estimates of rate constants are the same by both methods. The only advantage of the Bayesian approach seems to be in the estimation of uncertainties in cases where parameters are not well-defined by the data.

## Acknowledgments

I am very grateful to the following people for discussions and comments on drafts of this paper. Stephen Senn (Competence Center in Methodology and Statistics, CRP-Santé, Luxembourg). Tim Cole (UCL Institute of Child Health), Andrew Plested (Leibniz-Forschungsinstitut fur Molekulare Pharmakologie, Charite Universitatsmedizin, Berlin). Harvey Motulsky (CEO, GraphPad Software, Inc. La Jolla, USA) and Dr Rosi Sexton (Solihull, UK). And thanks to Dr Colin Longstaff (Biotherapeutics, NIBSC, South Mimms, EN6 3QG) for help with the web calculator…Thanks also to David Spiegelhalter (Cambridge, UK) for suggesting the term false positive risk (rather than false positive rate, that was used in earlier drafts).

Web calculator for false positive risk. This calculator does the calculations that are done by the following three R scripts. It works in any web browser (including those on smart phones) http://fpr-calc.ucl.ac.uk/

*Notes-on-use-of-R-scripts.pdf.* A brief description of how to run R scripts

*calc-prior.R* Calculates prior probability needed to produce specified FPR, printout and graphs of prior, P(H_1_), against observed *P* value. Sample output file: calc_prior-p=0.05-FPR=0.05.txt

*calc-pval.R* Calculates the *P* value that would be needed to achieve a specified false positive risk (given the prior probability, P(H_1_), and sample size). Sample output file:calc_pval for FPR=0.05 p(H_1_) =0.5.txt

*calc-FPR+LR.R* Calculates FPR and likelihood ratios for specified *P* value and sample size. No graphs, just printout file. Sample output file:calc-FPR+LR-p=0.05.txt

*Plot-FPR-vs-Pval.R* Generates plots of FPR as a function of observed *P* value. Plots are generated for 3 sample sizes (n =4, 8, 16 by default) and for two values of the prior, P(H_1_)=0.1 and 0.5 by default. Each plot is produced with arithmetic scales and log-log scales. Also, for direct comparison of the *p-equals* case and the *p-less-than* case, the FPR vs *P* value for each case are plotted on the same graph, for *n* = 16 (default), No print-out files in thus version.

*two_sample-simulation-+LR+prior.R* Simulates a specified number of *t* tests and prints out results (like in ref [1] but prints also the likelihood ratios and prior probabilities for “significant”; results. Plots graphs of the distributions from which random samples are generate, and the distribution of effect sizes and *P* values. Sample output file;: simulation+LR-P-between 0.0475 0.0525.txt

*t-distributions.R* Plots the distributions of Student’s *t* for the null and alternative hypothesis, for specified sample size and *P* value (defaults are *n* = 16 and value = 0.05). Used for Figure 1.

## References

1. Colquhoun D (2014) An investigation of the false discovery rate and the misinterpretation of p-values. R Soc Open Sci 1: 140216. 10.1098/rsos.140216[doi];rsos140216 [pii].

2. Berger JO, Sellke T (1987) Testing A Point Null Hypothesis - the Irreconcilability of P-Values and Evidence-Journal of the American Statistical Association 82: 112–122.

3. Berger JO, Delampady M (1987) Testing Precise Hypotheses. Statistical Science 2: 317–352.

4. Button KS, Ioannidis JP, Mokrysz C, Nosek BA, Flint J, Robinson ES, Munafo MR (2013) Power failure: why small sample size undermines the reliability of neuroscience. Nat Rev Neurosci 14: 365–376. nrn3475 [pii]; 10.1038/nrn3475 [doi].

5. Colquhoun D (2016) The problem with p-values. Aeon Magazine. Available: https://aeon.co/essays/it-s-time-for-science-to-abandon-the-term-statistically-significant

6. Berkson J (1942) Tests of Significance Considered as Evidence. Journal of the American Statistical Association 37: 325–335.

7. Sellke T, Bayarri MJ, Berger JO (2001) Calibration of p values for testing precise null hypotheses. American Statistician 55: 62–71.

8. Wacholder S, Chanock S, Garcia-Closas M, El GL, Rothman N (2004) Assessing the probability that a positive report is false: an approach for molecular epidemiology studies. J Natl Cancer Inst 96: 434–442.

9. Ioannidis JP (2005) Why most published research findings are false. PLoS Med 2: e124. Available: http://journals.plos.org/plosmedicine/article?id=10.1371/journal.pmed.0020124

10. Colquhoun D (2015) Response to comment by Loiselle and Ramchandra. Royal Society Open Science 2: 2150319. Available: http://rsos.royalsocietypublishing.org/content/royopensci/2/8/150319.full.pdf

11. Goodman SN (1993) p values, hypothesis tests, and likelihood: implications for epidemiology of a neglected historical debate. Am J Epidemiol 137: 485–496.

12. Goodman SN (1999) Toward evidence-based medical statistics. 1: The P value fallacy. Ann Intern Med 130: 995–1004. 199906150-00008 [pii].

13. Senn, S. J. (2007) Statistical Issues in Drug Deelopment. John Wiley & Sons, Ltd.

14. Colquhoun D Statistics and the law: the prosecutor’s fallacy. Available: http://www.dcscience.net/2016/03/22/statistics-and-the-law-the-prosecutors-fallacy/

15. Matthews RAJ (2001) Why should clinicians care about Bayesian methods? Journal of Statistical Planning and Inference 94: 43–58.

16. Wasserstein RL, Lazar NA (2017) The ASA’s Statement on p-Values: Context, Process, and Purpose. American Statistician 70: 129–133.

17. Royal Statistical Society ASA statement on P-values and statistical significance: Development and impact. Available: https://www.youtube.com/watch?v=B7mvbOK1ipA

18. Matthews R, Wasserstein R, Spiegelhalter D (2017) The ASA’s p-value statement, one year on. Significance 14: 38–41.

19. Heneghan C, Goldacre B, Mahtani KR (2017) Why clinical trial outcomes fail to translate into benefits for patients. Trials 18: 122. 10.1186/s13063-017-1870-2 Available: https://trialsjournal.biomedcentral.com/articles/10.1186/s13063-017-1870-2

20. Edwards, A. W. F. (1992) Likelihood. Expanded edition. The Johns Hopkins University Press.

21. Goodman SN (1999) Toward evidence-based medical statistics. 2: The Bayes factor. Ann Intern Med 130: 1005–1013. 199906150-00009 [pii].

22. Berger JO, Sellke T (1987) Testing A Point Null Hypothesis - the Irreconcilability of P-Values and Evidence. Journal of the American Statistical Association 82: 112–122.

23. Johnson VE (2013) Revised standards for statistical evidence. Proc Natl Acad Sci U S A 110: 19313–19317. 1313476110 [pii]; 10.1073/pnas.1313476110 [doi]. Available: http://www.pnas.org/content/110/48/19313.full.pdf?with-ds=yes

24. Curtis MJ Experimental design and analysis and their reporting: new guidance for publication in BJP. Available: http://onlinelibrary.wiley.com/doi/10.1111/bph.12856/full

25. Johnson VE (2013) UNIFORMLY MOST POWERFUL BAYESIAN TESTS. Annals of Statistics 41: 1716–1741. 10.1214/13-AOS1123 [doi].

26. Colquhoun, D. (2014) Two more cases of hype in glamour journals: magnets, cocoa and memory. Available: http://www.dcscience.net/2014/11/02/two-more-cases-of-hype-in-glamour-journals-magnets-cocoa-and-memory/<[28]

27. Open Science Collaboration (2015) Estimating the reproducibility of psychological science. Science 349: DOI: 10.1126/science.aac4716.

28. Kunert R Yet more evidence for questionable research practices in original studies of reproducibility projectt: psychology. Available: https://brainsidea.wordpress.com/2016/04/25/yet-more-evidence-for-questionable-research-practices-in-original-studies-of-reproducibility-project-psychology/

29. Colquhoun D (2007) How to get good science. Physiology News 69, 12–14 2007 Available at http://www.dcscience.net/2007/08/03/how-should-universities-be-run-to-get-the-best-out-of-people/.

30. Colquhoun D Publish and perish at Imperial College London: the death of Stefan Grimm. Available: http://www.dcscience.net/2014/12/01/publish-and-perish-at-imperial-college-london-the-death-of-stefan-grimm/

31. Goldstein H, Spiegelhalter DJ (1996) League Tables and Their Limitations: Statistical Issues in Comparisons of Institutional Performance. Journal of the Royal Statistical Society Series A 159: 385–443.

32. Varin C, Cattelan M, Firth D (2015) Statistical modelling of citation exchange between statistics journals. Journal of the Royal Statistical Society Series A 179: 1–33.

33. Curry, S. University rankings are fake news. How do we fix them? Available: http://occamstypewriter.org/scurry/2017/05/16/university-rankings-are-fake-news/

34. Senn, S. J. Double Jeopardy?: Judge Jeffreys Upholds the Law (sequel to the pathetic P-value). Available: https://errorstatistics.com/2015/05/09/stephen-senn-double-jeopardy-judge-jeffreys-upholds-the-law-guest-post/2017

35. Colquhoun D, Hawkes AG, Srodzinski K (1996) Joint distributions of apparent open times and shut times of single ion channels and the maximum likelihood fitting of mechanisms. Philosophical Transactions of the Royal Society London A 354: 2555–2590.

36. Colquhoun D, Hatton CJ, Hawkes AG (2003) The quality of maximum likelihood estimates of ion channel rate constants. J Physiol (Lond) 547: 699–728.

37. Epstein M, Calderhead B, Girolami MA, Sivilotti LG (2016) Bayesian Statistical Inference in Ion-Channel Models with Exact Missed Event Correction. Biophys J 111: 333–348. S0006-3495(16)30450-7 [pii]; 10.1016/j.bpj.2016.04.053 [doi].

38. Bullied Into Bad Science. Available: http://bulliedintobadscience.org/

39. Benjamin, D. and 71 others (2017). Redefine statistical significance. PysArXiv preprints, Available https://osf.io/preprints/psyarxiv/mky9j/ Accessed 24 July 2017.

40. Holder, R. (2009) The Bayesian interpretation of a P-value depends only weakly on statistical power in realistic situations Journal of Clinical Epidemiology 62, 1242–1247

41. Held, L. (2013). Reverse-Bayes analysis of two common misinterpretations of significance tests. Clinical Trials 2013 Apr;10(2):236–42. doi: 10.1177/1740774512468807

42. Colquhoun, D. (2015). There are powerful currents whipping up the metric tide. The HEFCE metrics report Availaible at http://www.dcscience.net/2015/07/09/there-are-powerful-currents-whipping-up-the-metric-tide-the-hefce-metrics-report/

43. Smaldino, P. E., and McElreath, R. (2016). The natural selection of bad science. Royal Society Open Science DOI:10.1098/rsos.160384 Available at http://rsos.royalsocietypublishing.org/content/3/9/160384

44. Computer programs used in this paper. Click here to download all R programs, and instruction for use in a zip file

